# The species problem from the modeler’s point of view

**DOI:** 10.1101/075580

**Authors:** Marc Manceau, Amaury Lambert

**Author notes:** Corresponding author Marc Manceau.

## Abstract

How to define a partition of individuals into species is a long-standing question called the *species problem* in systematics. Here, we focus on this problem in the thought experiment where individuals reproduce clonally and both the differentiation process and the population genealogies are explicitly known. We point out three desirable properties of species partitions: (A) Heterotypy between species, (B) Homotypy within species and (M) Monophyly of each species. We then ask: How and when is it possible to delineate species in a way satisfying these properties?

We point out that the three desirable properties cannot in general be satisfied simultaneously, but that any two of them can. We mathematically prove the existence of the finest partition satisfying (A) and (M) and the coarsest partition satisfying (B) and (M). For each of them, we propose a simple algorithm to build the associated phylogeny out of the genealogy.

The ways we propose to phrase the species problem shed new light on the interaction between the genealogical and phylogenetic scales in modeling work. The two definitions centered on the monophyly property can readily be used at a higher taxonomic level as well, e.g. to cluster species into monophyletic genera.

## Introduction

Models in macro-evolution have traditionally been centered on species. The so-called *lineage-based models* of diversification form a wide class of models considering species as key evolutionary units, thought of as particles that can give birth to other particles (i.e., speciate) during a given lifetime (i.e., before extinction) (see Stadler 2013; Pyron and Burbrink 2013; Morlon 2014 for reviews). In contrast, evolutionary processes amenable to direct empirical measurement (differentiation, reproduction, selection) are usually described at the level of individuals or populations.

The Neutral Theory of Biodiversity (NTB) (Hubbell 2001) opened a new way of thinking about species in macro-evolution. The birth, death, differentiation and speciation processes are described at the level of individuals, under the assumption of selective neutrality. In the last two decades, a popular way of studying macro-evolution has followed, consisting in performing computer-intensive simulations of individual-based stochastic processes of species diversification (Jabot and Chave 2009; Aguilée et al. 2011; Rosindell et al. 2015; Gascuel et al. 2015; Missa et al. 2016). These models rely on three major steps: (i) The genealogy of individuals is produced under a fixed or stochastic scenario of population dynamics; (ii) A process of phenotypic differentiation superimposed on the genealogy generates a partition of individuals into phenotypic groups; (iii) A species definition is postulated and is used to cluster individuals into different species, in relation to both the genealogy and phenotypic groups. These three steps allow modelers to track the evolutionary history of species, where extinction and speciation events emerge from the genealogical history of individual organisms.

The scenario of population dynamics (i) has for example been modeled with the Wright-Fisher and Moran model from population genetics or with density-dependent branching processes (Durrett 2008). We will not focus on that step, but we will nonetheless consider throughout the paper that individuals reproduce clonally. The genealogical relationships within a sample *χ* of present-day individuals will thus be a known rooted tree denoted *T*. Each tip of *T* is labelled by an element of *χ*, each internal vertex corresponds to some ancestor of elements of *χ*, and an edge between vertices represents a parent-child relationship. After running through step (ii), all individuals in *χ* can be grouped into clusters of individuals on the basis of their phenotype. We call the associated *χ*-partition the *phenotypic partition*, denoted 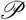.

In this paper, we review how steps (ii) and (iii) have been handled in the literature, before asking the following theoretical question:

> *How* and *when* is it possible to delineate species, in a way satisfying *biologically meaningful properties*, in an ideal situation where the phenotypic partition 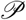 is specified and the entire genealogy *T* is known?

Because we work under the simplifying assumption of clonally reproducing organisms, this question is formally identical to the problem of defining and delineating *genera* when the *species phylogeny* is known, or any similar question formulated at a higher-order level (Aldous et al. 2008, 2011).

In the biological literature, the problem of agreeing on what should be considered as the most relevant concept of species is a long-standing question called the *species problem*. The species problem is both a conceptual question (defining the species concept) and a practical problem (classifying individuals into species) (Bock 2004). Several of the most notable evolutionary biologists (e.g. Dar-win, Dobzhansky, Mayr, Simpson, Hennig…) took a stand on the species problem often leading to vigorous debates (see Mayden 1997; De Queiroz 2007 for overviews of historical disagreements).

In the very simple setting that we are looking at, two classes of species concepts appear relevant in the quest for *biologically meaningful properties* of species. The ‘typological species concepts’ (Regan 1925; Sneath 1976) correspond to the clustering of individuals on the sole basis of their observed phenotype. This can be translated into two desirable properties: (A) any two individuals in distinct clusters can be distinguished based on at least one characteristic, (B) individuals belonging to the same cluster share the same characteristics. The later foundation and spread of cladistics by Hennig (Hennig 1965) marked a radical change of paradigm in the systematic classification, which quickly resulted in the proposition of so-called ‘phylogenetic species concepts’ (De Queiroz and Donoghue 1988; Avise and Ball 1990; Baum 2009). These definitions, which brought to the forefront the notion of common ancestry, provide us with a third desirable property: (M) species are monophyletic groups of individuals, i.e. any two individuals in one cluster are more closely related to each other than either is to any individual in another cluster.

Ideally, defining putative species in our framework amounts to finding clusters of individuals satisfying these three desirable properties that we will denote throughout the paper:

(A) Heterotypy between species
(B) Homotypy within species
(M) Monophyly

We quickly observe that by definition, the phenotypic partition 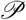 satisfies the two desirable properties: (A) and (B). Unfortunately, phenotypically similar individuals may not be more genealogically related to one another than to any different individual. In other words 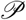 does not in general satisfy (M). Convergence and reversal events, by making a trait either appear several times independently in different parts of the tree or by making a trait disappear in subtrees, are classically invoked to explain the non-monophyly of 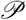. However, let us stress that even with traits evolving without convergence or reversal, individuals characterized by an ancestral phenotype may define a non-monophyletic subset of *χ*, a phenomenon called ‘ancestral type retention’ or ‘plesiomorphy’ (see Figure 2).

**Fig. 2:**
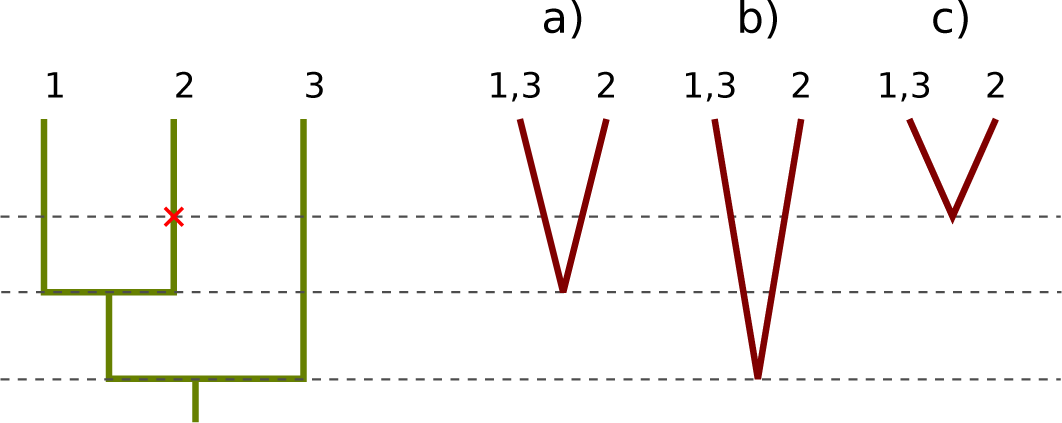
Building the phylogeny out of the genealogy. Left panel: A fixed genealogy with one mutational event giving rise to a derived character responsible for partitioning {1, 2, 3} into the two distinct phenotypic groups {1, 3} and {2}, assumed to be different species. Right panel: Phylogenies associated with three possible choices of divergence times, from left to right: a) Shortest or b) Longest coalescence time between individuals of different species; c) Date of origin of the derived character.

The paper is organized as follows. In Section 1, we review the main species definitions used in the context of individual-based modeling of macro-evolution. In Section 2, we introduce the formalism required to study species partitions, and we make some preliminary observations on the three desirable properties. In Section 3, we prove mathematically that it makes sense to define the finest species partition satisfying (A) and (M) and the coarsest species partition satisfying (B) and (M). We call these respectively the *loose* and the *lacy* species partitions. Finally, we discuss the relevance of these propositions from both empirical and theoretical points of view.

## 1 Five species definitions in individual-based models

We provide here an overview of five modes of speciation that have been proposed so far in the context of individual-based models of diversification (see Kopp 2010 for a review). Among these five modes, only the second one is intended to model specifically the geographical isolation of two subpopulations. The four other modes focus on modeling sympatric speciation by means of gradual accumulation of mutations. These five propositions are illustrated in Figure 1.

**Fig. 1:**
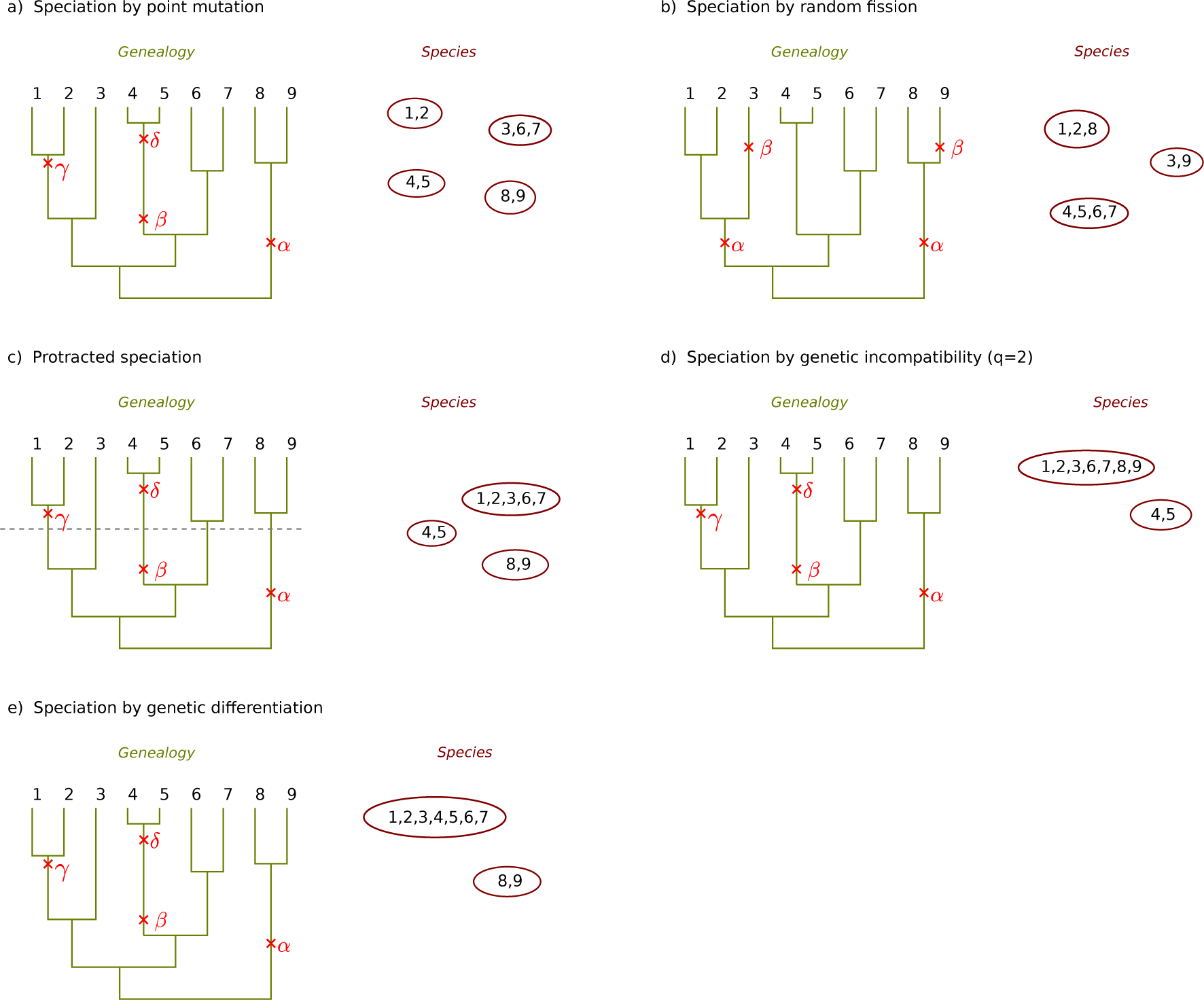
The five modes of speciation proposed in individual-based models of macro-evolution. In each panel, the genealogy (tree) of individuals (integer labels) is given on the left, along with mutations (crosses) that confer new types (Greek letters) to individuals. The corresponding species partition is represented on the right of each panel (subsets of labels circled).

### Speciation by point mutation

This mode of speciation was proposed in the original framework of the NTB (Hubbell 2001; Jabot and Chave 2009). Differentiation occurs as the product of neutral mutations modeled by a point process on the genealogy. Each mutation confers a new type to the lineage carrying it (infinite-allele model) and to its descent before any new mutation arises downstream. Species are then defined as groups of individuals carrying the same type. The phenotypic partition and the species partition thus coincide by definition.

### Speciation by random fission, or peripheral isolates

These two closely related models have also been proposed first in the framework of the NTB (Hubbell 2001, 2003), but see also Lambert and Ma (2015). In these models, independently of the genealogy, each phenotypic class of individuals, interpreted as a geographic deme, may split at random times into two new demes. In Figure 1b, this is illustrated as mutations hitting simultaneously several lineages in the same phenotypic class, which endows them with the same new phenotype. The two propositions differ only with regard to the size of the newly formed deme, whether the split is even (random fission) or uneven (peripheral isolate).

### Protracted speciation

This model intends to reflect the general idea that speciation is not instantaneous (Rosindell et al. 2010; Lambert et al. 2015; Etienne et al. 2014). The differentiation process is usually assumed to be differentiation by point mutation under the infinite-allele model. A new phenotypic class is called an incipient species but becomes a so-called good species only after a fixed or random time duration. In other words, two individuals belong to different species if they carry different phenotypes and if they diverged far enough in the past. In Figure 1c, the species arisen from the mutation labelled γ is still incipient at present time. More complex models of protracted speciation feature several stages that incipient species have to go through before becoming good species.

### Speciation by genetic incompatibility

This generalization of the point mutation mode of speciation (Melián et al. 2012) is inspired by the model of Bateson-Dobzhansky-Muller incompatibilities (Orr 1995). Again, a first step consists in endowing the genealogy of individuals with neutral mutations. Then two individuals are said compatible if there are fewer than *q* mutations on the genealogical path linking them. Finally, species are the connected components of the graph associated to the compatibility relationship between individuals. For *q* ≠ 1, there can be incompatible pairs of individuals in the same species, as can be seen in Figure 1d with individuals labelled 1 and 9 for example. Speciation by point mutation corresponds to the particular case *q* =1.

### Speciation by genetic differentiation

This model of speciation also assumes that phenotypic differentiation is driven by point mutations on the genealogy. Species are then defined as the smallest monophyletic groups of individuals such that any pair of individuals carrying the same phenotype are always in the same species (Manceau et al. 2015). We will show later that this definition, hereafter called *loose species definition*, always makes sense once given a phenotypic partition and a genealogy.

As can be seen in Figure 1, the first four models out of the five described in the previous section yield partitions of individuals into species that are in general non-monophyletic with respect to the underlying genealogy. This is problematic when it comes to measuring the phylogenetic relationship between species, reflecting their shared evolutionary history (Velasco 2008). In particular based on the true genealogy, there are multiple, arbitrary ways of defining the divergence time between two non-monophyletic species. Readers will convince themselves that none of the three solutions illustrated in Figure 2 gives a reasonable account of the genealogical history subtending species. As part of the current effort to bridging the gap between micro- and macro-evolution (Graham and Fine 2008; Rosindell et al. 2011; Pennell and Harmon 2013), it is crucial to interlock the genealogical and phylogenetic scales. This is the rationale behind considering monophyly as a desirable property of species constructions.

In Sections 2 and 3, we will consider as given the genealogical tree *T* of the set of all present-day organisms *χ*, and the partition 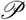 of *χ* into phenotypic groups. The tree *T* may have been generated under any model of population dynamics and the partition 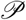 may have been produced by any process of differentiation unfolding through time. With these data at hand, we formalize the three desirable properties of the species partition mentioned in the introduction and then study different ways to fulfill them.

## 2 Three desirable properties of species definitions

For each internal node of *T*, by a slight abuse of terminology, we call *clade* the subset of *χ* comprising exactly all tips descending from this node. We denote by *ℋ* the collection of all clades of *T*. Note that as a subset of *χ*, *χ* itself is an element of *ℋ*, and that for every *x* ∈ *χ*, the singleton {x} is an element of *ℋ*. Moreover, any two clades *C* and *D* elements of *ℋ*, are always either nested or mutually exclusive, meaning that *C* ∩ *D* can only be equal to *C, D* or ø. Mathematically, a collection of nonempty subsets of *χ* satisfying these properties is called a *hierarchy*, and it can be shown that to any hierarchy corresponds a unique rooted tree with tips labelled by *χ*. Therefore, we will equivalently speak of *T* or of its hierarchy *ℋ*. For a nice discussion around the notion of hierarchy and neighboring concepts, see Steel (2014).

One should keep in mind that *ℋ* and 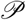 are both collections of subsets of *χ*, but that *ℋ* is not a partition. With this formalism, we define the species problem as: Given *ℋ* and 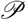 find a partition 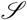 of *χ*, called the *species partition*, whose elements are called *species clusters* or simply *species*, satisfying one or more of the following three desirable properties:

A. *Heterotypy between species.* Individuals in different species are phenotypically different, i.e. for each phenotypic cluster 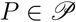 and for each species cluster 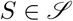, either *P* ⊆ *S* or *P* ∩ *S* = ∅;
B. *Homotypy within species.* Individuals in the same species are phenotypically identical, i.e. for each phenotypic cluster 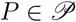 and for each species cluster 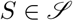, either *S* ⊆ *P* or *P* ∩ *S* = ∅.
C. *Monophyly.* Each species is a clade of *T*, i.e. 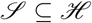 As mentioned in the introduction, if 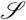 satisfies both (A) and (B), then it is immediate from the preceding definitions that 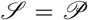. If in addition 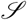 satisfies (M) then 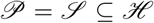. We record this as a first observation.

### Observation 1

*Unless we are given 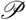 and ℋ such that 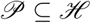(that is, each phenotypic cluster is a clade in the first place), no species partition satisfies simultaneously (A), (B) and (M)*.

Then our next question is: ‘Is there a species partition 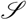 for which two of them hold?’ For X, Y equal to A, B or M, we will write (XY) the property (X AND Y).

We already saw that 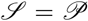 satisfies (AB). Now let us go for species partitions 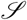 satisfying (M). To fulfill (A), each 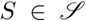 must contain all the phenotypic clusters it intersects. So in particular the partition 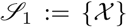 fulfills (AM). This trivial solution corresponds to assigning all the individuals of *χ* to one single species. Symmetrically, to fulfill (B), each 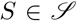 must be contained in all the phenotypic groups it intersects. So in particular the partition 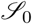 made of all singletons fulfills (BM). This trivial solution corresponds to assigning each individual of *χ* to a different species. This is recorded in the following observation.

### Observation 2

*For any 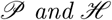 and *ℋ* and for any two desirable properties among (A), (B) and (M), there is at least one species partition 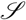 satisfying both properties*.

The species partitions 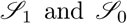 are obviously not biologically relevant. In particular, we would like to find species partitions that are *finer* than assigning all individuals to one single species, or *coarser* than assigning each individual to a different species.

We use the standard notions of finer and coarser partitions of a set (Bóna 2011). Let 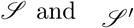 be two partitions of the set *χ*. We say that 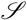 is finer than 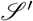, and we write 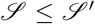 if for each 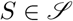 and each 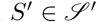, either *S* ⊆ *S′* or *S* ∩ *S′* = ∅. If 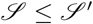, we say equivalently that 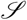 is finer than 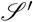 or that 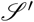 is coarser than 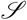.

Remark that two species partitions 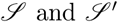 cannot always be compared, in the sense that they can satisfy neither 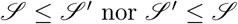. The relation ≤ is thus not a linear order on all the partitions of *χ*, but is known to be a partial order (see Appendix A for details).

Now observe that properties (A) and (B) can precisely be stated in terms of inequalities associated with the partial order ≤ as follows.

### Observation 3

*Consider a given phenotypic partition 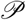 and a species partition 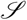.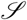 satisfies (A) if and only if 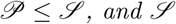 satisfies (B) if and only if 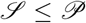. As a consequence, if 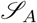 satisfies (A) and 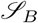 satisfies (B), then*

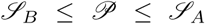

This leads us to investigate the possibility of defining ‘the finest partition satisfying (AM)’ as well as ‘the coarsest partition satisfying (BM)’.

## 3 The lacy and loose species definitions

In general, there is no guarantee that the coarsest or finest partition of a given set of partitions does belong to this set. However, we state the following result that ensures the existence of the finest partition satisfying (AM) and the coarsest partition satisfying (BM).

### Theorem 1

*Given 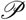 and ℋ, there exists a unique finest partition of *χ* satisfying (AM), and a unique coarsest partition of *χ* satisfying (BM)*.

This result is proved in Appendix B. It allows us to highlight and name two new different species definitions:

The *loose species definition* is the finest partition satisfying (AM).

The *lacy species definition* is the coarsest partition satisfying (BM).

For any species partition 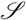 satisfying (M), there is a unique phylogenetic tree 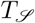 which represents the evolutionary relationships between the species in 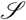 consistently with the genealogy *T*. For every species *S*, since *S* is monophyletic there is a unique internal node *u*(*S*) of *T* such that *S* is exactly constituted of the labels of the tips subtended by *u*(*S*). Then 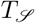 is obtained from T by merging, for every species S, the subtree descending from *u*(*S*) into a single edge. This is expressed in terms of hierarchies in the following observation.

Observation 4

*Consider a given genealogical hierarchy ℋ and species partition 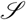 satisfying (M). The hierarchy 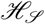 corresponding to the phylogenetic tree 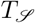 can be defined by*

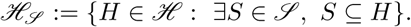

So for both the loose and the lacy species partition, there is a phylogeny consistent with the genealogy. Figure 3 shows both the lacy phylogeny and the loose phylogeny associated with a simple genealogy and a simple phenotypic partition.

**Fig. 3:**
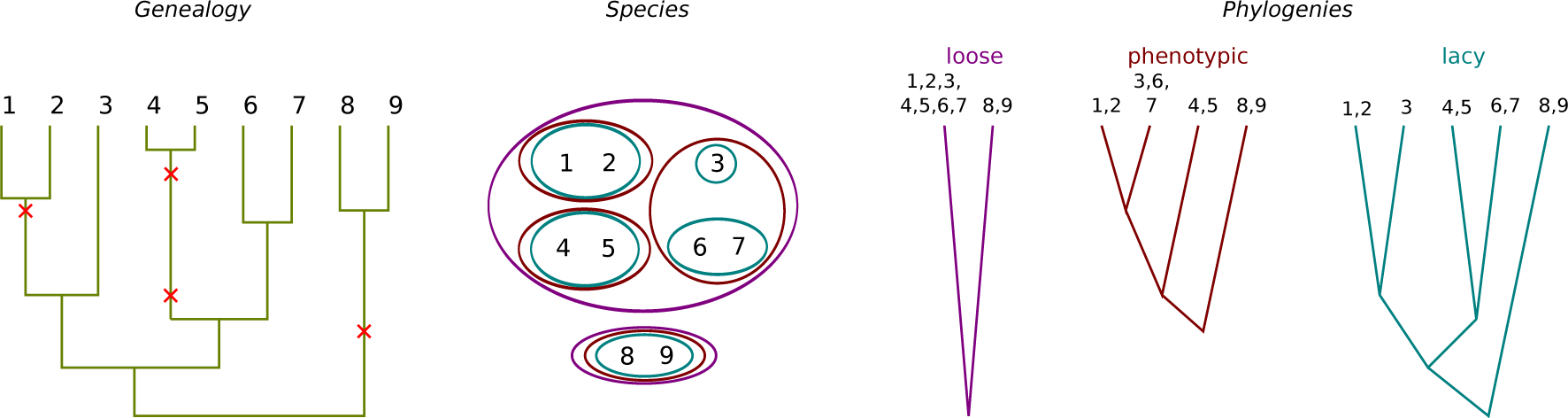
Species partitions associated to each of three definitions. Left panel: The fixed genealogy with point mutations (infinite-allele model) leading to the phenotypic partition 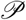 = {{1, 2}, {3, 6, 7}, {4, 5}, {8, 9}}. Middle panel: Inclusion relations between the three species partitions, as discussed in Observation 3, the loose partition is coarser than the phenotypic partition, which is coarser than the lacy partition. Right panel: Phylogenies corresponding to the three species partitions, from left to right: loose, phenotypic (under the arbitrary convention that divergence times are taken as mutation times), lacy.

For any genealogy *T* and phenotypic partition 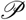, we now describe a procedure to get the phylogeny corresponding either to the lacy or to the loose definition, without requiring the knowledge of species partitions. Interestingly, building 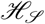 this way offers a quick way to get 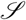 under the lacy and loose definitions, because species are the smallest sets of labels in 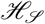. The different steps of the algorithm are explained hereafter, illustrated in Figure 4 and formalized in Appendix C.

**Fig. 4:**
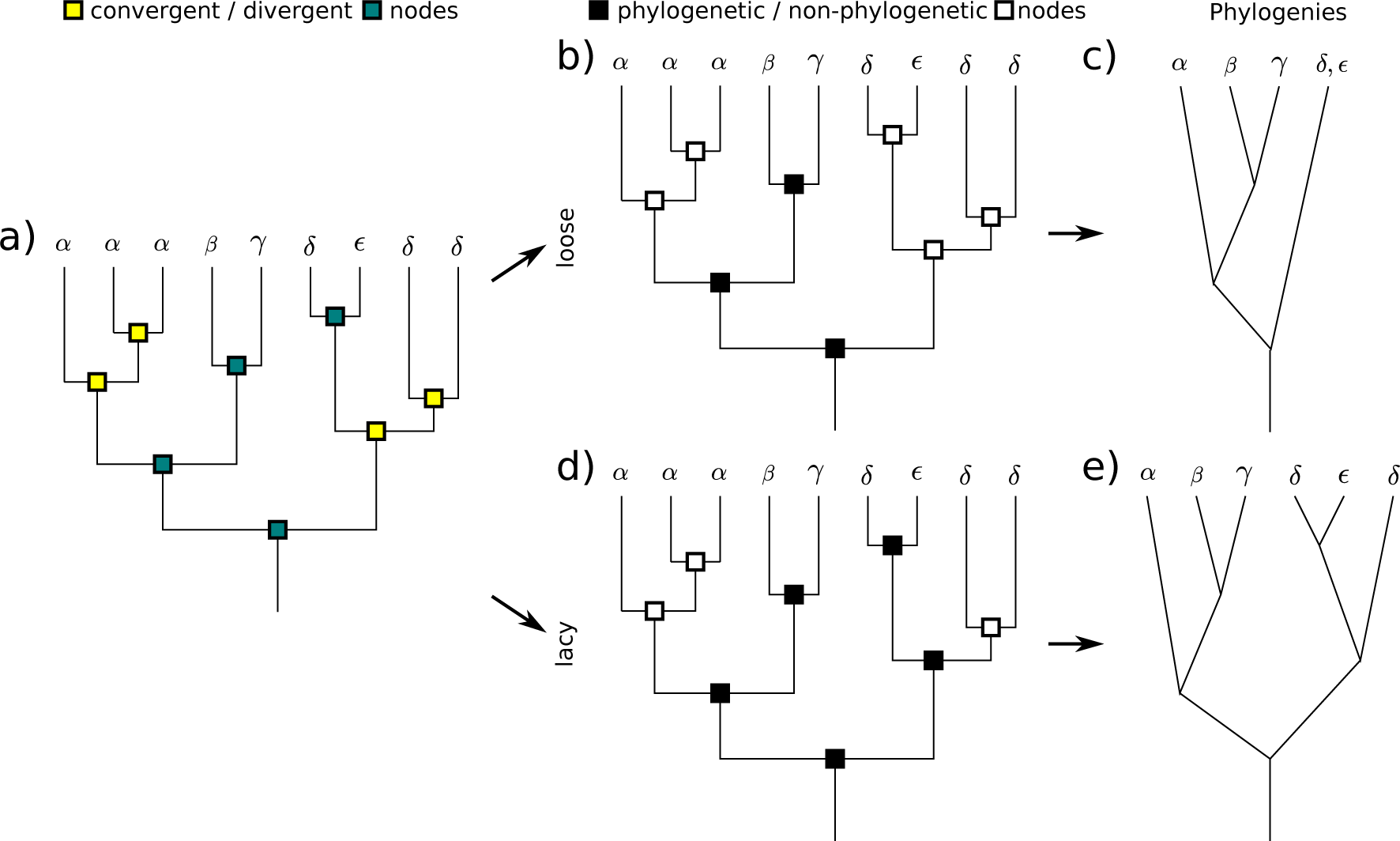
Construction of the phylogeny under the lacy and loose species definitions. Greek letters correspond to phenotypes. a) The genealogy with interior nodes classified as convergent (light yellow) or divergent (dark blue). bd) The genealogy with interior nodes classified as non-phylogenetic (white) or phylogenetic (black). ce) The corresponding phylogeny. b) Loose: Light nodes and dark nodes descending from light nodes are colored white. d) Lacy: Dark nodes and light nodes ancestors of a dark node are colored black.

First, we classify all interior nodes of the genealogy as *convergent node* or *divergent node*. An interior node is *convergent* if there are at least two tips, one in each of its two descending subtrees, carrying the same phenotype. Otherwise the node is said to be *divergent*. Note that convergent nodes may be ancestors of divergent nodes when the phenotypic partition is not monophyletic. Second, we build a phylogeny by deciding which interior nodes are *phylogenetic nodes*, that is, appear in the phylogeny.

### Observation 5

*The loose phylogeny is obtained by declaring non-phylogenetic (i) all convergent nodes and (ii) all divergent nodes descending from a convergent node. Other nodes are declared phylogenetic*.

*The lacy phylogeny is obtained by declaring phylogenetic (i) all divergent nodes and (ii) all convergent nodes ancestral to divergent nodes. Other nodes are declared non-phylogenetic*.

The last observation holds due to the following reasons:

By definition, the two clades *C* and *C′* subtended by a convergent node satisfy *C* ∩ *P* ≠ ∅ and *C′* ∩ *P* ≠ ∅ for some phenotypic cluster 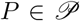. As a consequence, these two clades have to be included in the same species cluster in a species partition satisfying the heterotypy property (A), that is, a convergent node cannot appear in a phylogeny satisfying (A). Conversely, any phylogeny whose nodes are included in the set of divergent nodes of the genealogy satisfies (A). The finest partition satisfying (A) corresponds to the phylogeny containing the largest number of divergent nodes, and only divergent nodes, as in the construction of the loose phylogeny proposed in the observation.

Symmetrically, for the two clades *C*,*C′* subtended by a divergent node, we have that *C* ∩ *P* ≠ ∅ implies *C′* ∩ *P* ≠ ∅ for any phenotypic cluster 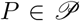. As a consequence, these two clades have to belong to two different species clusters in a species partition satisfying the homotypy property (B), that is, any divergent node has to appear in a phylogeny satisfying (B). Conversely, any phylogeny whose nodes contain all divergent nodes of the genealogy satisfies (B). The coarsest partition satisfying (B) corresponds to the phylogeny containing the smallest number of convergent nodes, but all divergent nodes, as in the construction of the lacy phylogeny proposed in the observation.

## Discussion

The present study builds on recent attempts to describe evolutionary trees on various scales simultaneously. The multi-species coalescent is one of the most influential of these attempts (Maddison 1997). In this model, a species tree is first specified and given the species tree, the gene genealogies are drawn from a censored coalescent (i.e., ancestral lineages can coalesce only if they lie in the same ancestral species). In particular, this approach has been used to assess the relevance of the reciprocal monophyly criterion to recognize species (Hudson and Coyne 2002; Mehta et al. 2016). Note that this framework introduces a top-down coupling between the macro-evolutionary scale and the micro-evolutionary scale, thus relying on an external species definition. In contrast, the bottom-up approach that we adopted consists in assuming that macro-evolutionary patterns are shaped by micro-evolutionary processes. Let us point out two studies similar in spirit to ours, which make several proposals to lump together lower-order taxa (e.g. species) in order to build trees on higher-order taxa (e.g. genera) (Aldous et al. 2008, 2011). The authors ground their definitions on the knowledge of a phylogeny with point mutations, the main differences being that each branching event distinguishes a mother and a daughter lineage, and that monophyly is not a desirable property in their work.

On the contrary, we put to the forefront the monophyly property (M), together with (A) heterotypy between species and (B) homotypy within species. We explicitly defined and compared three species definitions, each of these satisfying a different set of properties: Phenotypic (AB), Loose (AM) and Lacy (BM).

Additionally, we stress that the most popular way of defining species in individual-based modeling studies of macro-evolution (i.e., speciation by point mutation) in general leads to non-monophyletic species. In contrast, the loose species definition, previously used in the context of macro-evolution (Manceau et al. 2015), systematically yields monophyletic species. Here we extended this study and compared it to a third species definition also satisfying (M), the lacy species definition. Finally, we provided a standardized procedure to build the lacy and loose species partitions given a genealogy and a phenotypic partition.

In practice, the task of systematists is the inference of ancestral relationships between individual organisms from molecular sequence and phenotype data and the characterization of species from those data. Classifying diversity is notoriously difficult for many reasons, including the difficulty of choosing the appropriate level of description, the ubiquitous presence of convergent evolution and reversal events, and the difficulty to agree on a unique species concept (Mayden 1997; De Queiroz 2007; Baum 2009).

On the other hand, the task of modelers, assuming a fully known individual-based evolutionary history, appears at first sight trivial. They face, however, the same difficulty in defining a proper species concept. Even within the very simple framework that we considered, three distinct species definitions came out. They all fit the general species definition of *‘separately evolving metapopulation lineages’* (De Queiroz 2007), while satisfying distinct desirable properties. We argue that comparing species definitions based on the properties they fulfill in simple models might help shed light on the species problem.

These definitions, despite their simplicity, might also provide a coarse-grained picture of the way different real-world taxa are described. For example, species described long ago are more likely to be based on phenotypic information alone, and seem thus closer to the phenotypic species partition. Conversely, the rise of molecular methods in evolutionary biology may have brought (M) to the forefront. The reconstruction of gene genealogies has stimulated the development of methods aiming at automatically delineating putative species from sequence data: from a single phylogeny (Fujisawa and Barraclough 2013), from gene trees (Yang and Rannala 2010), or from raw molecular data (Puillandre et al. 2012). Recent species descriptions are thus more likely to concern monophyletic groups of individuals than earlier. The lacy definition could be used as crude picture of recent taxonomist work on emblematic clades. In these clades, more or less homotypic groups of individuals can be separated on a genealogical basis only into what are known as *cryptic species* (Bickford et al. 2007). In other clades, taxonomists may prefer to ensure that species are diagnosable and monophyletic units. These two properties, stressed as ‘priority taxon naming criteria’ (Vences et al. 2013), are closer in agreement with the loose definition.

Note that advanced theoretical work has been undertaken in a framework closer to biological reality, but much less connected to most modeling studies in macro-evolution (Dress et al. 2010; Kwok 2011; Alexander 2013; Alexander et al. 2015). While we based our study on the knowledge of only one genealogy, even the genealogical history of supposedly ‘asexual’ real-world organisms such as bacteria shows evidence for horizontal gene transfer events (Puigbò et al. 2013). The genealogical history of organisms should in general be represented as a non-tree network, or as a collection of gene genealogies, making far more complex the question of grouping individuals into taxa (Hudson and Coyne 2002; Samadi and Barberousse 2006).

Individual-based modeling is a promising avenue for understanding macro-evolution from first principles, as it may allow evolutionary biologists to describe explicitly the stochastic demography of whole metacommunities and the ecological interactions between different types of individuals in each community. We believe that these processes may have left enough signal in both the shape of evolutionary trees and the patterns of contemporary biodiversity, so as to be unraveled by statistical inference. Understanding how species, the elementary units of macro-evolution, are formed and deformed by these processes remains a major challenge, to which the present work hopefully contributes.

## Acknowledgements

The authors are very grateful to F. Débarre, R.S. Etienne, M. Steel, S. Türpitz and A. Hoppe for their comments on this paper, and to D. Baum for helpful literature advice. The authors thank the Center for Interdisciplinary Research in Biology (CIRB, Collège de France) for funding, as well as the École Normale Supérieure for MM PhD funding. We declare no conflict of interest.

## Appendix The species problem from the modeler’s point of view

Some of the results stated in Sections A and B are classical results in combinatorics for partially ordered sets (see Bóna 2011, chapter 16). For the sake of self-containment and because all readers may not be familiar with these notions, we nevertheless expose them here.

### A ‘Finer than’, a partial order relation on *χ*-partitions

#### Definition 1

Let 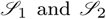 be two *χ*-partitions. We say that 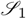 is finer than 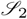, and we write 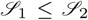 if 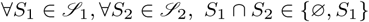.

We detail here the three criteria that make the ‘finer than’ relation a partial order on the set of *χ*-partitions.

*Proof* One must check the reflexivity, antisymmetry and transitivity properties.

- Reflexivity. Take any *χ*-partition 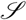. Then for all 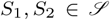 we either have *S*_1_ ∩ *S*_2_ = *S*_1_ if *S*_1_ = *S*_2_, or *S*_1_ ∩ *S*_2_ = ∅ otherwise. It follows that 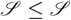.
- Antisymmetry. Take two *χ*-partitions denoted 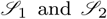, verifying 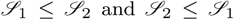. Then for all 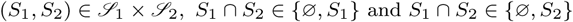. If *S*_1_ ∩ *S*_2_ ≠ ∅, it follows that *S*_1_ = *S*_2_, and finally 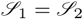.
- Transitivity. Take now three *χ*-partitions denoted 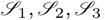, verifying 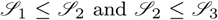. Let 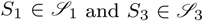 and assume that *S*_1_ ∩ *S*_3_ ≠ ∅. Then there is *x* ∈ *S*_1_ ∩ *S*_3_ and we let *S*_2_ be the unique element of 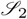 such that *x* ∈ *S*_2_. Thus *S*_1_ ∩ *S*_2_ ≠ ∅ and *S*_2_ ∩ *S*_3_ ≠ ∅, which implies by assumption that *S*_2_ ∩ *S*_1_ = *S*_1_ and *S*_2_ ∩ *S*_3_ = *S*_2_. So we see that *S*_1_ ⊆ *S*_2_ ⊆ *S*_3_, so that *S*_1_ ∩ *S*_3_ = *S*_1_.

### B Proof of Theorem 1

Here we will consider sets of partitions verifying one or two desirable properties. Hence the following definitions

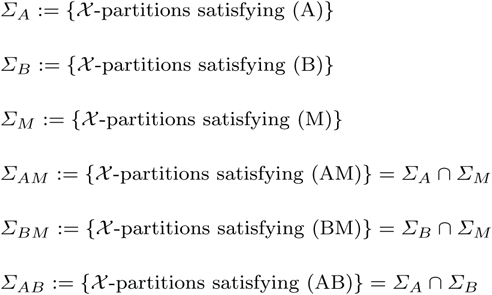

We will see that the collection of *χ*-partitions *∑_M_* plays a singular role in Theorem 1. This is due to the characterization of *∑_M_* by the fact that there is a hierarchy *ℋ* (here the hierarchy associated with the genealogy *T*) such that

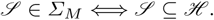

Also recall that the collections of *χ*-partitions *∑_A_* and *∑_B_* can be defined as follows

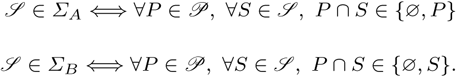

In this section, we aim at giving a proof of Theorem 1 which can now be restated as follows

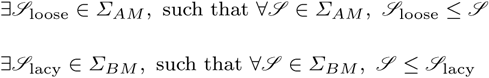

The proof is divided into two parts. First, given a set of partitions ∑ (resp. ∑ ⊆ *∑_M_*), we prove the existence of the finest (resp. coarsest) partition finer (resp. coarser) than any element of ∑, which we call inf ∑ (resp. sup ∑). Second, we show that inf *∑_AM_* ∈ *∑_AM_* and sup *∑_BM_* ∈ *∑_BM_*, hence yielding the definitions 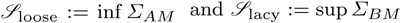.

#### B.1 Defining the supremum and the infimum of a set of *χ*-partitions

##### Definition 2

For any non-empty collection ∑ of *χ*-partitions, we define the two relations 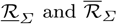 and on *χ* by

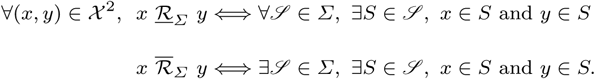

##### Lemma 1

*For any non-empty collection ∑ of *χ*-partitions, 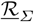 is an equivalence relation. For any non-empty collection ∑ of *χ*-partitions such that ∑ ⊆ ∑_M_, 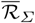 is an equivalence relation*.

*Proof* The *reflexivity* and *symmetry* of the two relations are easily seen. Now let us prove their transitivity. Let ∑ be a non-empty collection of *χ*-partitions, and (*x,y,z*) ∈ *χ*^3^ such that 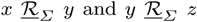. Let 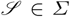. By definition,

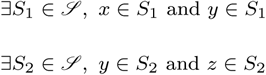

It follows that *y* ∈ *S*_1_ ∩ *S*_2_, and because 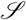 is a partition, *S*_1_ = *S*_2_. Finally, with *S* := *S*_1_ = *S*_2_, there exists 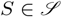 such that *x* ∈ *S* and *z* ∈ *S*, so that 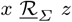 and we can conclude that 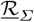 is transitive.

Now let ∑ ⊆ ∑_M_ be a non-empty collection of *χ*-partitions and (*x,y,z*) ∈ *χ*^3^ such that 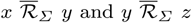.

By definition,

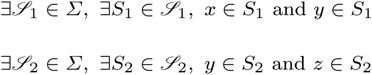

Because ∑ ⊆ ∑_M_, 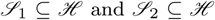, so that 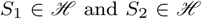. From the definition of hierarchy, we get *S*_1_ ∩ *S*_2_ ∈ {∅, *S*_1_, *S*_2_}. Since *y* ∈ *S*_1_ ∩ *S*_2_, we have *S*_1_ ∩ *S*_2_ ≠ Ø

Suppose that *S*_1_ ∩ *S*_2_ = *S*_2_. It follows that 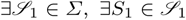, *x* ∈ *S*_1_ and *z* ∈ *S*_1_.

Suppose that *S*_1_ ∩ *S*_2_ = *S*_1_. It follows that 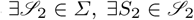, *x* ∈ *S*_2_ and *z* ∈ *S*_2_. So 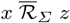 and we can conclude that 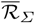 is transitive.

##### Definition 3

For any non-empty collection ∑ of *χ*-partitions, we call inf ∑ the *χ*-partition induced by the equivalence relation 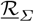. For any non-empty collection ∑ of *χ*-partitions such that ∑ ⊆ ∑_M_, we call sup ∑ the *χ*-partition induced by the equivalence relation 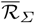.

Readers familiar with lattice theory will note that these definitions match the usual ‘meet’ and ‘join’ operators used for lattices, and in particular the lattice of partitions of a set, ordered by refinement. For the other readers, recall first that any equivalence relation on a set *χ* induces an *χ*-partition obtained by placing all elements in relation in one cluster. Further, the following lemma justifies the notation inf and sup.

##### Lemma 2

*Let ∑ be any non-empty collection of *χ*-partitions. Then for any 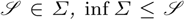. Let ∑ be any non-empty collection of *χ*-partitions such that ∑ ⊆ ∑_M_. Then for any 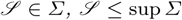*.

##### Proof

Let ∑ be any non-empty collection of *χ*-partitions and *S* ∈ inf ∑. Let also 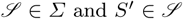. We need to prove that *S* ∩ *S′* ∈ {∅, *S*}. Assume that *S* ∩ *S′* ≡ ∅ and *S* ∩ *S′* ≠ *S*. Then there is *x* ∈ *S* ∩ *S′* and *y* ∈ *S* such that *y* ∉ *S′*. Because *x,y* ∈ *S*, by definition of inf ∑, we have 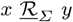 and by definition of 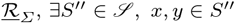. So *S′* and *S″* are both elements of 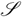 containing *x*, which implies that *S′* = *S″* and contradicts *y* ∉ *S′*.

Now let ∑ be any non-empty collection of *χ*-partitions such that ∑ ⊆ ∑_M_ and *S* ∈ sup ∑. Let also 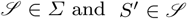. We need to prove that *S* ∩ *S′* ∈ {∅, *S*}. Assume that *S* ∩ *S′* ≡ ∅ and *S* ∩ *S′* ≡ *S*′ Then there is *x* ∈ *S* ∩ *S′* and *y* ∈ *S* such that *y* ∉ *S′*. Because *x* ∈ *S* and *y* ∉ *S*, by definition of sup *∑*, *x* and *y* are not in relation 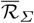 and by definition of 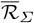, either *x* ∉ *S′* or *y* ∉ *S′* and we get a contradiction.

Note that, in general, we can have inf *∑* ∉ *∑* and sup *∑* ∉ *∑*. Here are two examples to provide the reader with some intuition.

*Example 1.* Take

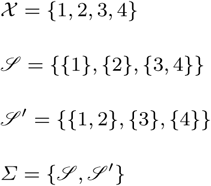

In this case, we get inf *∑* = {{1}, {2}, {3}, {4}}, which does not belong to *∑*. Moreover, if we define the hierarchy *ℋ* := {{1, 2, 3, 4}, {1, 2}, {3, 4}, {1}, {2}, {3}, {4}}, we have *∑* ⊆ *∑*_M_ which allows us to consider sup *∑* = {{1, 2}, {3, 4}} which again does not belong to *∑*.

*Example* 2.Take

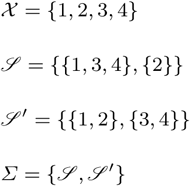

In this case, we get inf *∑* = {{1}, {2}, {3, 4}}, which does not belong to *∑*. Moreover, there is no *χ*-hierarchy *ℋ* such that 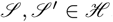. Then we can see that the relation 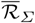 is not an equivalence relation on *χ*, because 1 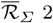 and 1 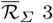, but we do not have 2 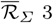. Thus, sup Σ is not defined.

#### B.2 Proving that inf **∑*_AM_* ∈ **∑*_AM_* and sup **∑*_BM_* ∈ **∑*_BM_*

In order to prove that inf **∑*_AM_* ∈ **∑*_AM_* and sup **∑*_BM_* ∈ **∑*_BM_*, we will rely on properties of inf *∑* and sup *∑* presented in the following lemma.

##### Lemma 3

*For any non-empty collection *∑* of *χ*-partitions, for any *S* ∈ inf *∑*, *S* can be written in the form of the following non-empty intersection*

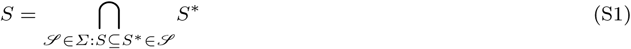

*For any non-empty collection *∑* of *χ*-partitions such that *∑* ⊆ *∑*_M_, for any *S* ∈ sup *∑*, *S* can be written in the form of the following non-empty union*

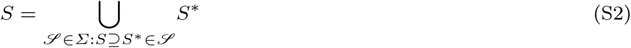

*In addition,*

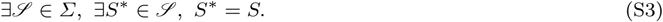

##### Proof

We begin with proving (S1). Let *∑* be any non-empty collection of *χ*-partitions and consider *S* ∈ inf *∑*. Now set

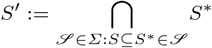

and let us prove that *S* = *S′*. According to Lemma 2 that for any 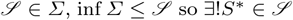, such that *S* ⊆ *S\**. This proves that the intersection in the definition of *S′* is not empty. Now by definition of *S′* we have *S* ⊆ *S′*, which also implies *S′* = ∅. We need to show now that *S* ⊆ *S′*. Let *x* be any element of *S′* and *y* be any element of *S*. Then for any 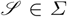, there is (a unique) 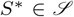 such that *S* ⊆ *S\** and by definition of *S′*, we have *x* ∈ *S\**. But since *S* ⊆ *S\** we also have *y* ∈ *S\**. This shows that for any 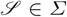 there is 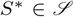 such that *x* ∈ *S\** and *y* ∈ *S\**. This can be expressed equivalently as 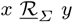, so that *x* and *y* are in the same element of inf *∑*, that is *x* ∈ *S*.

Now let us prove (S2). Let *∑* be any non-empty collection of *χ*-partitions such that *∑* ⊆ *∑*_M_ and let *S* ∈ sup *∑*.

Set

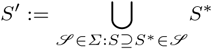

and let us prove that *S* = *S′*. According to Lemma 2 that for all 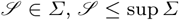, so 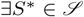 such that *S\** ⊆ *S*. In particular, the intersection in the definition of *S′* is not empty and *S′* = ∅. Now by definition of *S′* we have *S′* ⊆ *S*. We need to show now that *S* ⊆ *S′*. Let *x* be any element of *S* and *y* be any element of *S′*. Since *S′* ⊆ *S*, *y* ∈ *S* so that *x* and *y* are in the same element of sup *∑*, which can be expressed equivalently as 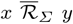. Now by definition of 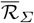, there is 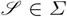 and 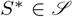 such that *x,y* ∈ *S\**. Now since *S\** ∩ *S* ≠ ∅, we have *S\** ⊆ *S*, which shows by definition of *S′* that *x* ∈ *S′*.

It remains to show (S3)

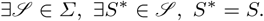

Let us prove by induction on *n* ≥ 1 that for any *F* ⊆ *S* of cardinality *n*, there is 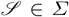 and 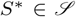 such that *F* ⊆ *S\** ⊆ *S*. The result will follow by taking *F* = *S*. For *n* =1, the property holds due to to (S2). Let *n* ≥ 1 strictly smaller than the cardinality of *S* and assume that the property holds for all integers smaller than or equal to *n*. Let *F* be any subset of *S* of cardinality *n* + 1 and write *F* = *F*_1_ ∪ {*x*}, where *x* ∉ *F*_1_. Since *F*_1_ is of cardinality *n* there is 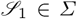 and 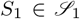 such that *F*_1_ ⊆ *S*_1_ ⊆ *S*. Let *y* ∈ *F*_1_. There is also 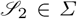 and 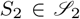 such that {*x,y*} ⊆ *S*_2_ ⊆ *S*. Now because 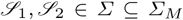, we have 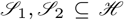, so that 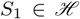 and 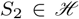. From the definition of hierarchy, we get *S*_1_ ∩ *S*_2_ ∈ {∅, *S*_1_, *S*_2_}. Since *y* ∈ *S*_1_ ∩ *S*_2_, we have *S*_1_ ∩ *S*_2_ ≠ ∅, so one of the two, denoted *S\** contains the other one. In particular, *F*_1_ ⊆ *S\** ⊆ *S* and {*x,y*} ⊆ *S\** ⊆ *S*, which shows that *F* = *F*_1_ ∪ {*x*} ⊆ *S\** ⊆ *S* and terminates the proof.

We can now go back to the proof of Theorem 1.

##### Proof

(i) inf **∑*_AM_* ∈ **∑*_A_*: Consider *S* ∈ inf **∑*_AM_* and 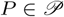. From Lemma 3, we get

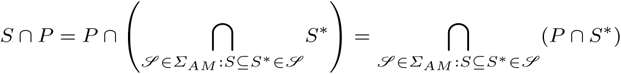

Now for each 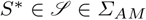, *P* ∩ *S\** ∈ {∅, *P*}, thus leading to *S* ∩ *P* ∈ {∅, *P*}, that is inf **∑*_AM_* ∈ **∑*_A_*.

(ii) inf**∑*_AM_* ∈ **∑*_M_*: Consider *S* ∈ inf **∑*_AM_*. From Lemma 3, we get

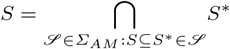

Now for each 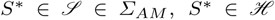. Moreover, the hierarchy *ℋ* is closed under finite, non-disjoint intersections, thus leading to *S* ∈ *ℋ*, that is inf **∑*_AM_* ∈ **∑*_M_*.

(iii) sup **∑*_BM_* ∈ **∑*_B_*: Consider *S* ∈ sup **∑*_BM_* and recall from Lemma 3 that there is 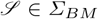 and 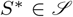 such that *S* = *S\**. Now for any 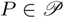,

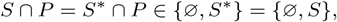

so that sup **∑*_BM_* ∈ **∑*_B_*.

(iv) sup **∑*_AM_* ∈ **∑*_M_*: Consider *S* ∈ sup **∑*_BM_* and *S\** = *S* as previously. Since *S\** ∈ *ℋ*, *S* ∈ *ℋ*, so that sup **∑*_BM_* ∈ **∑*_M_*.

This shows that inf **∑*_AM_* ∈ **∑*_AM_* and sup **∑*_BM_* ∈ **∑*_BM_*, which completes the proof of Theorem 1.

### C Construction of the lacy and loose phylogenies

This section aims at formalizing mathematically the construction of the lacy and loose phylogenies presented in the main text.

Recall that an interior node is convergent if there are two tips, one in each of its two descending subtrees, carrying the same phenotype, otherwise this node is said to be divergent. We will say that the two clades subtended by a convergent (resp. divergent) node are convergent (resp. divergent). We define *ℋ_d_* as the collection of divergent clades, that is

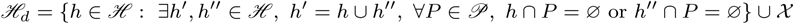

We similarly consider phylogenetic and non-phylogenetic clades for either the loose or the lacy definition. We call *ℋ*_loose_ and *ℋ*_lacy_ the collection of phylogenetic clades for the loose and lacy definitions respectively. The procedure described in the main text amounts to defining

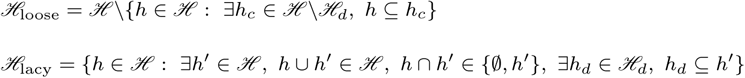

